# Targeted nanocarriers coopting pulmonary leukocytes for drug delivery to the injured brain

**DOI:** 10.1101/2022.02.04.479150

**Authors:** Patrick M. Glassman, Jia Nong, Jacob W. Myerson, Viviana Zuluaga-Ramirez, Alba Rodriguez-Garcia, Alvin Mukalel, Serena Omo-Lamai, Landis R. Walsh, Raisa Y. Kiseleva, Carlos H. Villa, Colin F. Greineder, Scott E. Kasner, Drew Weissman, Michael J. Mitchell, Silvia Muro, Yuri Persidsky, Jacob S. Brenner, Vladimir R. Muzykantov, Oscar A. Marcos-Contreras

## Abstract

Selective drug delivery to injured regions of the brain is an elusive, but biomedically important, goal. It is tempting to co-opt migrating white blood cells (WBC) to carry drugs to the injured brain, using natural WBC tropism. Current approaches to load cargoes to WBC have limited utility, particularly in acute conditions, due to the need for time consuming *ex vivo* manipulation and loading of cells. Physiological, *in vivo* loading of WBC may be advantageous in this scenario. Here we devised such a strategy, capitalizing on the unique features of the direct blood exchange between brain and lungs. Mediators emanating from the injured brain directly travel to the pulmonary vasculature via venous flow. In response to these mediators, WBCs, transiently residing in the pulmonary microvascular lumen, disembark and flow with arterial blood to the brain microvasculature, where they adhere and transmigrate to the brain parenchyma via the local chemoattractant gradient. We posited that direct *in vivo* targeting of cargoes to the pulmonary WBC pool may provide drug transfer to brain via this natural mechanism. To test this, we intravenously injected agents targeted to intercellular adhesion molecule 1 (ICAM) in mice with acute brain inflammation caused by direct injection of tumor necrosis factor alpha (TNF-α). We found that: A) At 2 hours, >20% of ICAM/NP accumulated in lungs, predominantly in WBCs; B) At 6 and 22 hours, ICAM/NP pulmonary uptake markedly decreased; C) In contrast, ICAM/NP uptake in brain increased ~5-fold in this time interval, concomitantly with migration of WBCs to the brain. Cranial window fluorescent microscopy confirmed WBC transport of ICAM/NP to the brain in TNF-α-challenged mice beyond the BBB. Importantly, demonstrating the pharmacologic relevance of this strategy, dexamethasone-loaded ICAM/liposomes abrogated brain edema in this model. In sum, coopting the natural homing of WBC from the lungs via ICAM-targeting to injured brain is an attractive strategy for precise interventions for treatment of acute brain injuries.

**VISUAL ABSTRACT:** 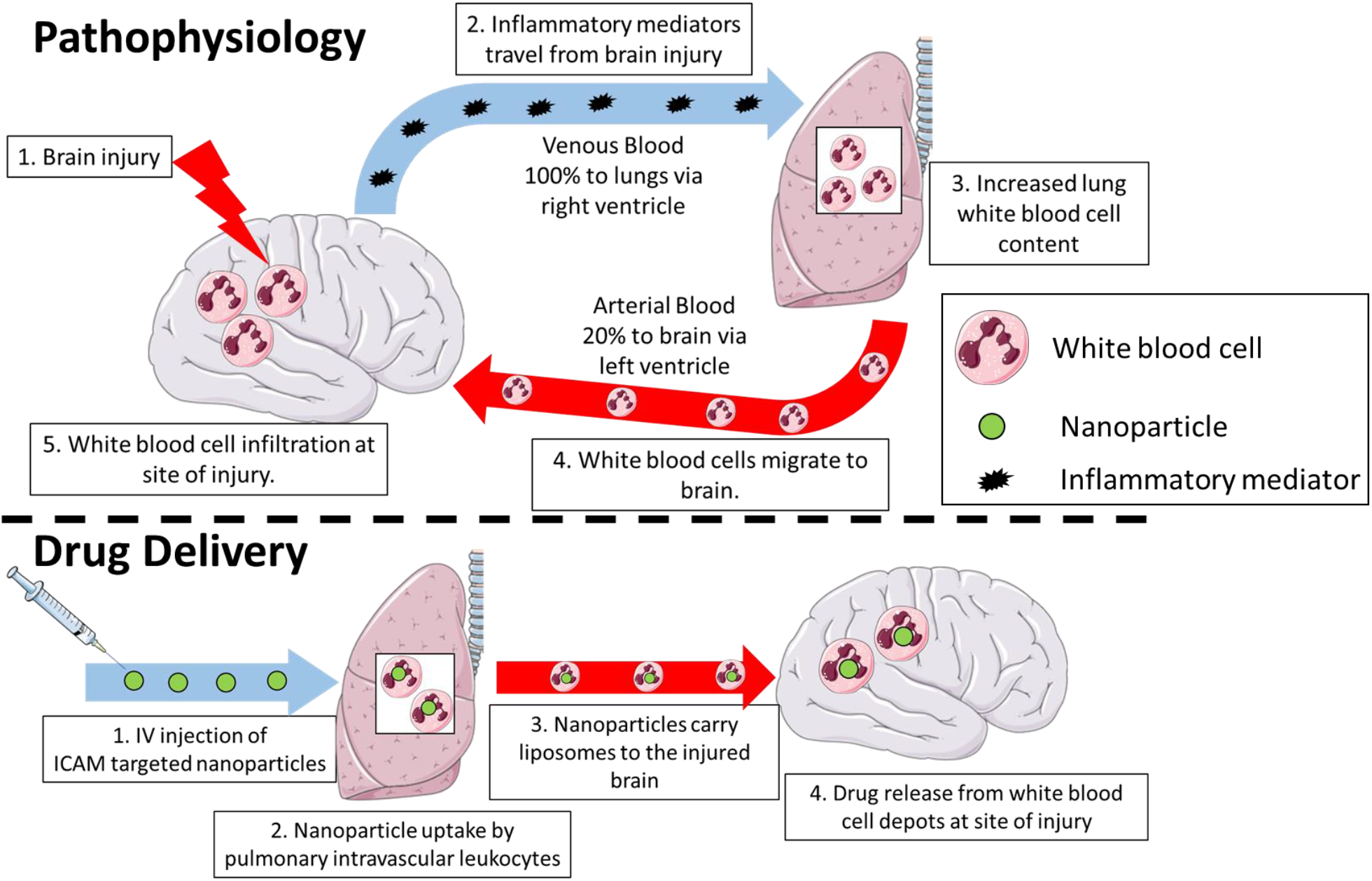

## INTRODUCTION

Targeted drug delivery to the brain promises breakthroughs in treatment of debilitating and lethal pathologies, including stroke, traumatic brain injury, glioblastoma and other brain tumors, meningitis, and neurodegenerative diseases^1, 2^. Various carriers with distinct chemistry, geometry, mechanical flexibility, and affinity have been devised to achieve this elusive goal^3, 4, 5, 6, 7^. One approach to enhance delivery employs targeting to and across the cerebral vasculature using antibodies, peptides, and other ligands of molecules that are stably expressed on the luminal surface of brain vessels. However, targeting to these molecules, including receptors for transferrin, insulin, and growth factors does not provide selectivity for sites of injury and inflammation^8, 9^. In order to achieve enhanced specificity for injured regions of the brain, targeting to inducible cell adhesion molecules (CAMs) expressed on endothelial cells, such as vascular CAM (VCAM)^9^, has been tested and has shown improved delivery and pharmacologic effects. Despite these inroads, direct, specific delivery to the parenchyma of the injured region of the brain remains an elusive goal.

To achieve the formidable goal of delivering drugs to the brain parenchyma within injured areas of the brain, it is tempting to utilize the natural homing of leukocytes to the pathologically altered region of the brain^10, 11, 12, 13^. This natural tropism is mediated by several processes including: A) activation of endothelial cells and white blood cells (WBC) by inflammatory mediators, B) increased exposure of adhesion molecules on both endothelial cells and WBC, C) disruption of endothelial tight junctions^14, 15^, and D) attraction of circulating leukocytes via a chemokine gradient emanating from the site of injury^16^. The idea of using isolated, *ex vivo* loaded WBC is attractive and several groups have reported therapeutic benefits of injecting drug-loaded WBCs in animal models of neurological disorders^17, 18, 19, 20, 21, 22^. Data on the fate of injected WBC (clearance, distribution, and effects on the body) are largely unknown. This aspect of the brain drug delivery requires systematic direct tracing of isotope-labeled agents^23^. However, *a priori*, this strategy is only permissive of loading a small number of cells and *ex vivo* manipulation may lead to undesired activation or alteration of these cells, resulting in severe adverse effects^24, 25, 26, 27^. Additionally, *ex vivo* manipulation of cells is not suitable for acute, emergency conditions and would require a specialized facility (e.g. as in CAR-T therapy) that is not likely to be found at most hospitals. A more desirable approach would be to specifically load those WBC that are predisposed to localize to the injured brain with drugs or drug carriers *in vivo*, bypassing the need for any *ex vivo* manipulation. It has been reported that ICAM is expressed on the surface of many WBCs, including monocytes and neutrophils^28^. Following inflammatory stimuli, the surface expression of ICAM on immune cells is significantly upregulated^29, 30, 31, 32^, providing selectivity for delivery to activated WBCs by targeting to ICAM.

### We postulate the ideal cells for this role are pulmonary intravascular leukocytes

The pulmonary circulation hosts the largest and most dynamic pool of intravascular WBC that are poised to quickly respond to local and remote signals from damaged tissues^33, 34^. There are no intervening capillary beds between the directly interconnected cerebral and pulmonary vasculatures. Hence, the constituent cells of the pulmonary vasculature (endothelial, pulmonary intravascular leukocytes) is the first set of extra-cerebral cells receiving signals from inflammatory mediators emanating from brain injuries (e.g. cytokines, exosomes, damage-associated molecular patterns). In fact, distal injuries often induce a secondary pulmonary pathology^35, 36, 37, 38^, which results in an increase in and hyperactivation of the pulmonary WBC pool. There are several reports that host defense cells responding to chronic neurological disorders mature in the lungs prior to trafficking to the injured brain^39, 40, 41^. These considerations imply that the dynamic pool of pulmonary WBCs are ideally positioned to shuttle drugs directly from the lungs to sites of brain pathology. However, strategies for controllable, specific, effective, and safe loading of drugs into intravascular WBCs have not been reported.

In the current study, we characterized the dynamic localization of leukocytes and ICAM-targeted pharmacological agents in the blood, lungs, and brain in a murine model of acute neurovascular inflammation induced by direct injection of tumor necrosis factor alpha (TNF-α) into the brain parenchyma. Our data presented below indicate that in the lungs, the number of WBC, the uptake of ICAM-targeted agents, and their fraction taken by intravascular WBC rapidly increased to the peak at 2 hours, followed by profound decline by 24 h. In the brain, in contrast, these cells gradually accumulated and increased ~5-fold 24 hours after TNF-α injury. Intravital fluorescent microscopy showed that in mice challenged with cranial injection of TNF, the migrating ICAM/NP-carrying WBCs accumulated in the brain parenchyma.

Leukocyte-mediated delivery to the brain parenchmya originating from ICAM-targeted lung uptake was demonstrated in the present study for monoclonal antibodies (mAb) and for three types of nanoparticles: polystyrene nanoparticles, liposomes, and lipid nanoparticles (LNP). IV injection of ICAM-targeted liposomes loaded with dexamethasone completely abrogated brain edema induced by TNF-α. These results indicate: A) <u>Direct, *in vivo* leukocyte loading:</u> after IV injection in mice with acute neurovascular inflammation, ICAM-targeted nanoparticles rapidly bind to pulmonary leukocytes *in vivo;* B) <u>Natural leukocyte trafficking:</u> these loaded leukocytes traffic to the inflamed region of the brain; and C) <u>Leukocyte-mediated drug delivery to the parenchyma:</u> this approach enables shuttling of nanoparticles to the site of brain injury, ultimately resulting in therapeutic efficacy. Overall, direct, *in vivo* targeting of the pulmonary WBC pool shortly after brain injury provides a mechanism to harness this dynamic pool of cells for selective drug delivery to the brain.

## RESULTS

### A systemic response to acute neurovascular inflammation in mouse model of intracranial TNF-α injection

In order to assess changes in accessible ICAM following brain injection of TNF-α, the tissue uptake of anti-ICAM (αICAM) mAb was investigated at several time points post injury. The direct quantitative measurements using isotope-labeled agents showed that: A) there was no significant differences in blood concentrations at different time points (**Figure 1a, Supplemental Table 1**); B) lung uptake reached a peak 2 hours post-injury and declined to baseline levels by 6 hours post-injury, suggesting a transient increase in ICAM levels in lungs post-brain injury (**Figure 1b**); C) brain uptake increased progressively with time after TNF-α insult, with αICAM brain delivery increasing 7-fold over naïve levels at 24 hours after injury (**Figure 1c**). Similar experiments were carried out for αCD45, which behaved with identical dynamics as αICAM, with a 4.6-fold increase in lung delivery 2 hours post-injury and a 16-fold increase in brain delivery 24 hours after injury. (**Figure 1d, e, f**).

**Figure 1.**
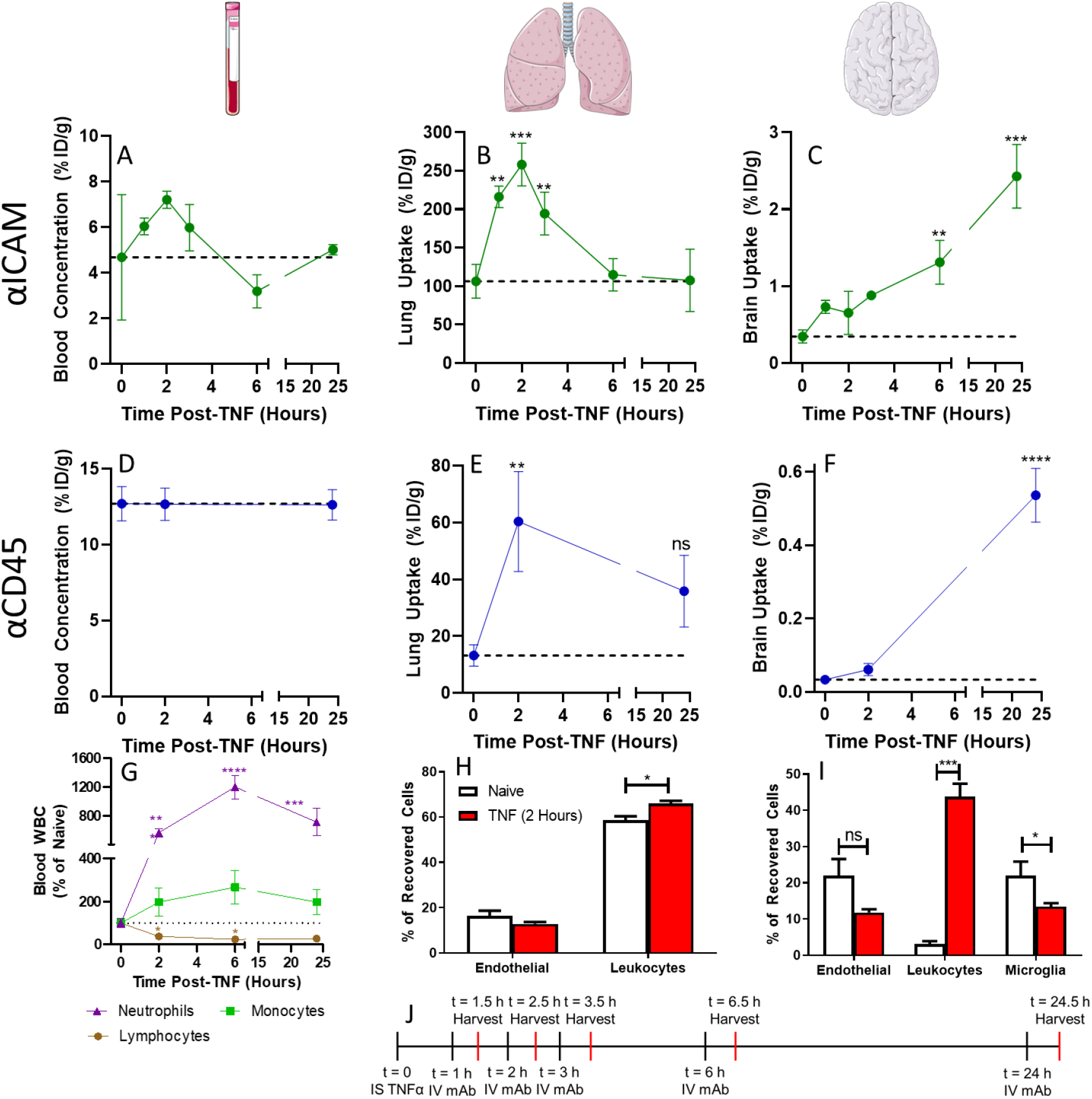
Local injection of TNF-α in the brain induces a systemic response. Following IV injection of αICAM, A) blood, B) lung, and C) brain targeting was assessed at several time points post-TNF. Similar studies were performed for αCD45 biodistribution in D) blood, E) lungs, and F) brain. Data represented as percent of injected dose per gram tissue (%ID/g). G) Complete blood counts were used to measure dynamic changes of white blood cells in circulation following TNF-α. Flow cytometry of single cell suspensions obtained from H) lungs 2 hours post-TNF-α and I) brain 24 hours post-TNF-α. Endothelial cells: CD31^+^CD45^-^, Leukocytes: CD45^+^, Microglia: CD45^mid^. J) Timeline of biodistribution experiments. Data represented as mean ± SEM. Dashed lines represent levels in naïve mice. Comparisons in A-G made by 1-way ANOVA with Dunnett’s post-hoc test vs. naïve mice and comparisons in H-I made by unpaired t-test. N=3/group.

Experiments were performed to evaluate dynamics of immune cells in blood, lungs, and brain following brain injury. Complete blood counts (CBC) revealed that the TNF-α injury affected circulating immune cells in several ways: A) transient increases in circulating neutrophils and monocytes, peaking 6 hours post-TNF-α; and B) a transient decrease in circulating lymphocytes, reaching a nadir at the same time point (**Figure 1g**). Flow cytometry evaluated cell type distributions in the lungs 2 hours after TNF and in the brain 24 hours after TNF, i.e., at the post-injury time points with the maximal level of αICAM uptake in the two corresponding organs. This analysis revealed that comparing with basal levels measured in naïve mice, at 2 hours post-injury, there was a significant increase in lung leukocytes (**Figure 1h**). 24 hours post-injury, there was a 14-fold increase in the relative recovery of brain leukocytes (**Figure 1i**).

These results are consistent with the following hypothetical spatiotemporal characteristics of the bi-directional vascular transport between the brain and lungs, illustrated in figure 2a. Pro-inflammatory mediators emanating from the site of brain injury are transported by blood pumped via the right heart chambers directly to the lungs collecting 100% of the venous blood ejected by the right ventricle. The mediators activate endothelial and white blood cells in the pulmonary vasculature, and ensuing interaction of these cellular constituencies further attract and activate circulating WBC to the lung microvasculature, serving as a transient “training base”, from which primed WBCs get transferred to the target organ passively with arterial blood flow (note: brain takes disproportionally high 15-20% of the cardiac arterial blood output), adhere to pathologically activated cerebral endothelial cells and transmigrate to the injured parenchyma. Furthermore, our results indicate that targeting to ICAM enables loading to the pulmonary WBCs permitting the subsequent trip to the brain just described above.

**Figure 2.**
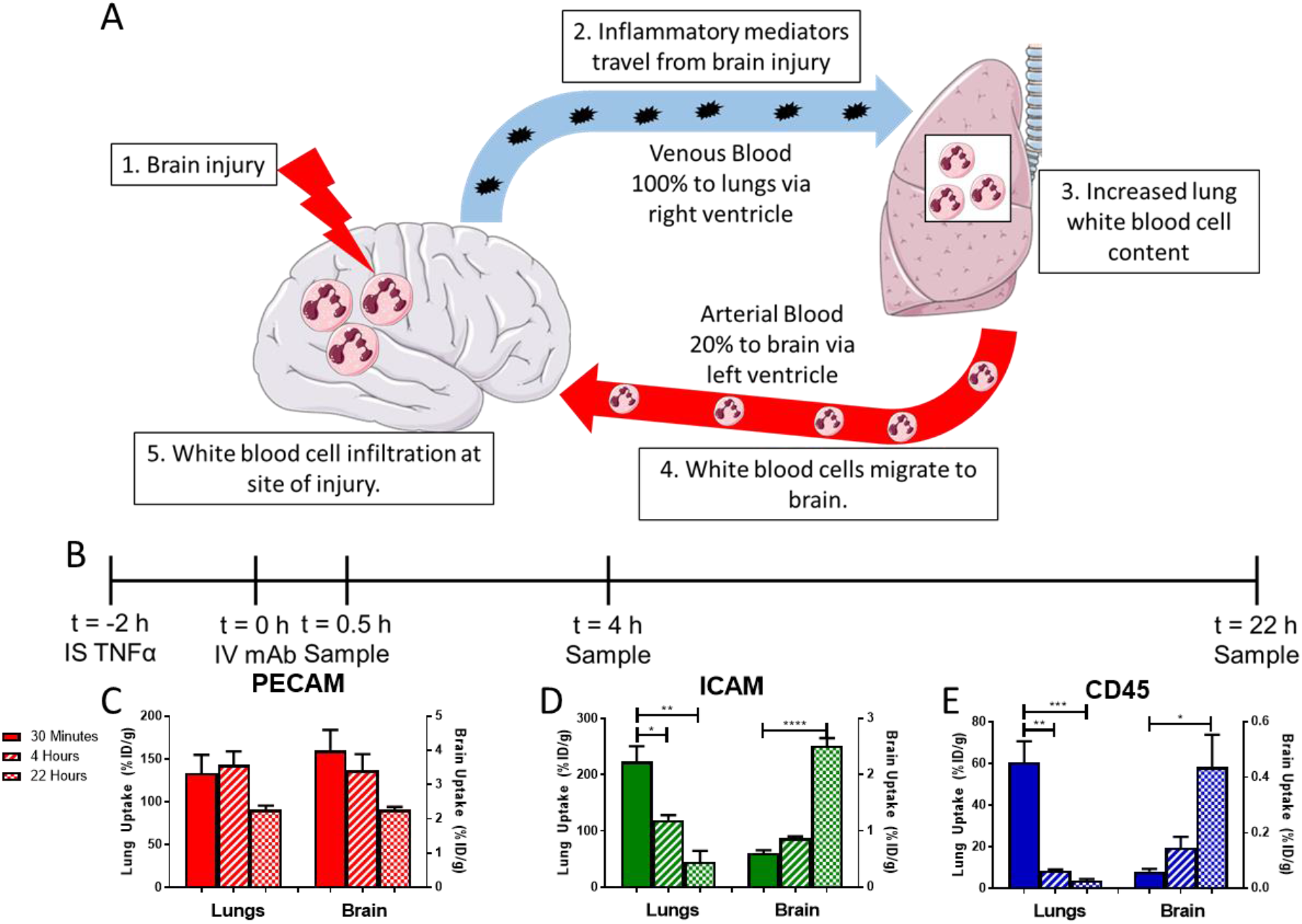
αICAM and αCD45 mAbs accumulate in the lungs, then migrate to the brain. A) Schematic of proposed mechanism underlying leukocyte migration. B) PK study timeline. Lung and brain pharmacokinetics of mAbs directed against: c) PECAM, D) ICAM, and E) CD45 following IV injection 2 hours post-TNF-α injury. Time points reflect the time post-mAb injection when organs were harvested. Data represented as mean ± SEM. Comparisons made by 1-way ANOVA with Dunnett’s post-hoc test vs. 30 minutes. N=3/group.

### ICAM-targeted monoclonal antibodies (mAbs) migrate to the brain

Encouraged by the identification of a lung-brain axis following brain injury (**Figure 2a**), we performed studies to appraise the utility of this novel drug delivery paradigm. Here, we injected isotope-labeled affinity ligands including αICAM into mice 2 hours post-TNF-α injury to evaluate the role of target epitope/cell type on pharmacokinetics and biodistribution (**Figure 2b**).

αPECAM behaved as expected for ligands of epitopes constitutively and stably on the surface of endothelial cells showing: A) specific (vs. IgG, see below) uptake in most organs at early time points; B) decreasing tissue concentrations over time (**Figure 2c**, **Supplemental Table 2**); and, C) more rapid blood clearance vs. control IgG (**Supplemental Figure 1**). In part due to more prolonged circulation time, control IgG slowly accumulated in the brain due enhanced vascular permeability, which has been previously reported in this model^9^ (**Supplemental Table 2, Supplemental Figure 2**).

The PK/BD of αICAM was more complex and rather unanticipated in some aspects. Over time, lung concentrations of αICAM decreased with a simultaneous <u>increase</u> in brain uptake αICAM (**Figure 2d, Supplemental Table 2**). The distribution pattern of αCD45 was similar to that of αICAM, with specific accumulation in lungs at early time points, followed by lung clearance and slow delivery to the brain (**Figure 2e, Supplemental Table 2**). Because CD45 is a pan-leukocyte marker, its accumulation can be attributed to an influx of mAb-tagged leukocytes at the injury in the brain. There was a significant correlation between clearance from the lung and changes in brain uptake with time (**Supplemental Figure 3**). It was hypothesized that this unexpected distribution pattern of αICAM was due to initial delivery of αICAM to activated leukocytes in the pulmonary vasculature followed by migration of leukocytes to the injured brain.

### Diversification of ICAM-directed loading of nanoparticles to lung WBC for subsequent delivery to the brain

For this purpose, we compared three different types of ICAM-targeted nanoparticles: polystyrene nanoparticles, liposomes, and LNP (**Figure 3a**). ICAM-targeted nanoparticles were largely cleared from the blood within 30 minutes; however, there was a rebound in blood concentrations over the next several hours for ICAM-targeted nanoparticles, potentially reflecting redistribution of leukocytes carrying nanoparticles into blood (**Supplemental Figure 4**). Similar to αICAM mAb, ICAM-targeted nanoparticles were largely taken up in the lungs within 30 minutes of injection (polystyrene nanoparticles: 147 ± 1 %ID/g, liposome: 174 ± 6 %ID/g, LNP: 123 ± 9 %ID/g), followed by clearance from the lungs over several hours (**Figure 3b, c, d, Supplemental Tables 3, 4, 5**). Both polystyrene nanoparticles and LNP displayed monotonic increases in brain concentrations with time after injection, while liposomes had a transient increase in brain uptake (**Figure 3b, c, d, Supplemental Table 4**). To evaluate the interplay between lung clearance and brain uptake of nanoparticles, lung/brain ratios were calculated at time points post-dose. All three particles displayed a steady increase in this ratio with time, reflecting the opposite trends in tissue targeting kinetics (**Figure 3e, f, g, Supplemental Tables 3, 4, 5**). On the contrary, untargeted control IgG nanoparticles did not display significant accumulation in either lungs or brain (concentrations > 10-fold lower than ICAM-targeted) (**Supplemental Figure 5, Supplemental Tables 3, 4, 5**).

**Figure 3:**
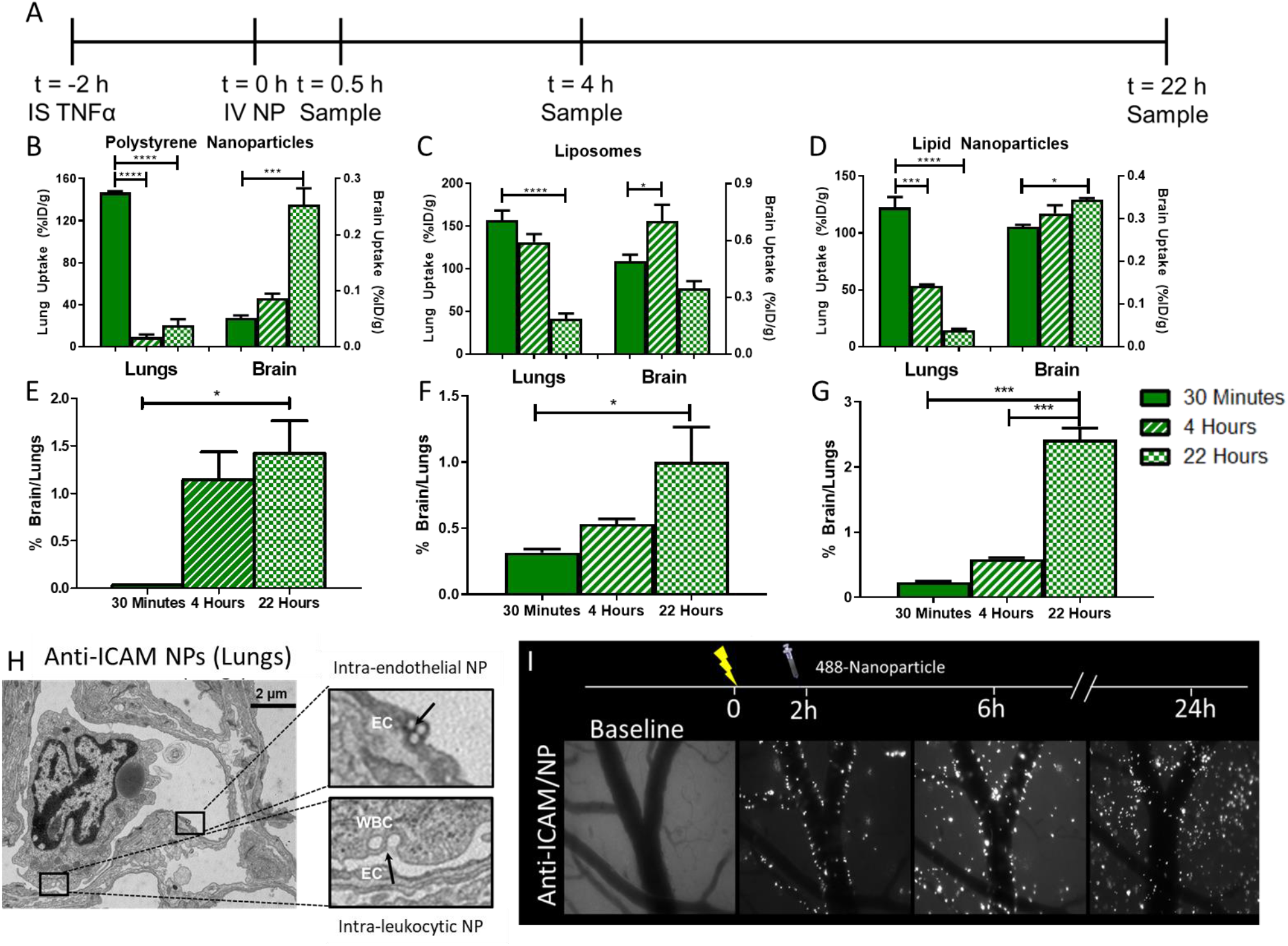
ICAM-targeted nanoparticles accumulate in the inflamed brain. A) Study timeline. Pharmacokinetics of B) polystyrene nanoparticles, C) liposomes, and D) lipid nanoparticles in lungs and brain following injection. Kinetic changes in the ratio of nanoparticles in brain vs. lungs for E) polystyrene nanoparticles, F) liposomes, and G) lipid nanoparticles. H) Transmission electron microscopy of ICAM-targeted polystyrene nanoparticles in lung endothelium and leukocytes 30 minutes post-injection. I) Cranial window intravital microscopy of ICAM-targeted polystyrene nanoparticles in TNF-α injured brain. Data represented as mean ± SEM. Comparisons in B, C, D were made by 1-way ANOVA with Dunnett’s post-hoc test vs. 30 minutes and those in E, F, G were made by 1-way ANOVA with Tukey’s post-hoc test. N ≥ 3/group.

Additional studies focused on visualizing the delivery mechanisms of ICAM-targeted nanoparticles in both lungs and brain. Transmission electron microscopy (TEM) demonstrated ICAM-targeted polystyrene nanoparticle localization to both endothelial cells and leukocytes in the lungs 30 minutes after IV injection (**Figure 3h**). Cranial window intravital microscopy (**Figure 3i**) showed; A) ICAM-targeted polystyrene nanoparticles were associated with the walls of inflamed brain blood vessels immediately following IV injection; B) consistent with radiotracing experiments, the number of nanoparticles in the cranial window increased over time after injection; C) 4 hours after injection, nanoparticles appeared in clusters and some beads were detected in the parenchyma; D) 22 hours after injection, nanoparticle fluorescence was no longer confined to large vessel walls and had spread into the parenchyma, suggesting that ICAM-targeted nanoparticles access a mechanism to cross the blood-brain barrier. Similar data were obtained for ICAM-targeted liposomes using cranial window intravital microscopy, with liposome fluorescence lining the vessel walls immediately post-injection and gradually accumulating in the brain parenchyma over 22 hours **(Supplemental Figure 6**). The fluorescent signal for liposomes was more diffuse than that for polystyrene nanoparticles, possibly reflecting differences in particle stability following internalization.

### ICAM targeted nanoparticles are predominantly delivered to leukocytes

Single cell suspensions were prepared from lungs 30 minutes after injection of ICAM-targeted nanoparticles (2 hours post TNF-α injury) (**Figure 4a**). Flow cytometry analysis showed that nearly all nanoparticle-positive cells in the lungs were leukocytes (CD45^+^) (93.4 ± 1.4% of recovered cells), with the remaining NC-positive cells being identified as endothelial cells (CD31^+^) (**Figure 4b**).

**Figure 4:**
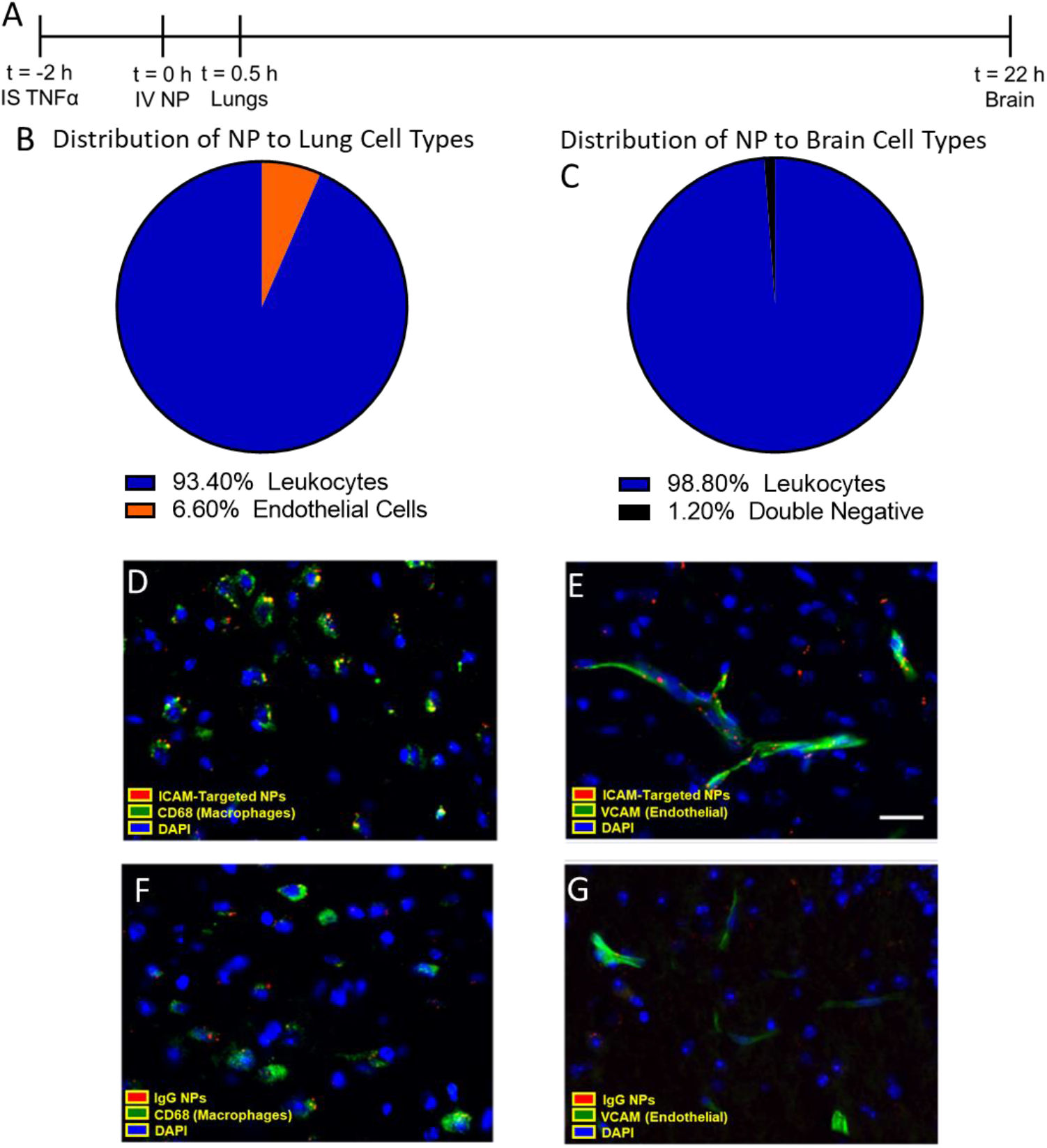
Cellular specificity of ICAM-targeted polystyrene nanoparticles. A) Flow cytometry was performed on single cell suspensions obtained from lungs and brain at the designated times post-nanoparticle injection. The fraction of nanoparticles recovered in B) lungs and C) brain that were associated with specific cell types. Leukocytes: CD45^+^, Endothelium: CD31^+^CD45^-^. Histology of brain tissue sections collected 22 hours post-injection of polystyrene nanoparticles in TNF-α challenged mice. Nanoparticle association with macrophages (CD68^+^) and endothelial cells (VCAM^+^) was measured for D, E) ICAM-targeted and F, G) IgG nanoparticles. Scale bar: 50 μm. Data represented as mean ± SEM. N = 3/group.

Having identified leukocytes as the primary target cells for ICAM-targeted nanoparticles in the lungs of TNF-α-challenged mice, we tested the hypothesis that these mobile leukocytes deliver αICAM/nanoparticles to the inflamed brain 22 hours after nanoparticle injection (24 hours post-injury). In single cell suspensions prepared from the brain, essentially all nanoparticle-positive cells were leukocytes (98.7 ± 0.2% of recovered cells) (**Figure 4c**). Flow cytometry showed polystyrene nanoparticle uptake in the brain for pristine and non-specific IgG-coated polystyrene nanoparticles, agreeing with biodistribution data (**Supplemental Figure 7**). A sub-typing of cells in the brain revealed that the majority of nanoparticle-positive leukocytes in the brain were monocytes/macrophages (73.0 ± 9.7%), with the bulk of the remainder being neutrophils (24.5 ± 9.9%) (**Supplemental Figure 8 and 9**). A large fraction of monocytes/macrophages were nanoparticle-positive (40.5 ± 3.6%). Among other leukocytes, 27.4 ± 6.6% of neutrophils and 25.2 ± 1.2% of other myeloid cells were nanoparticle-positive. Minimal ICAM-targeted nanoparticle uptake in microglia and T-cells (**Supplemental Figure 9b**).

Brain histology confirmed nanoparticle association with macrophages (CD68-stained) (**Figure 4d**, **Supplemental Figure 10a**) and endothelial cells (VCAM-stained) (**Figure 4e**, **Supplemental Figure 10b**). Histology indicated greater uptake of ICAM-targeted nanoparticles vs. IgG-coated nanoparticles, both in the vasculature and in the brain parenchyma (**Figure 4f, g, Supplemental Figure 10a, b**). Parenchymal nanoparticle fluorescence was largely co-localized with macrophages, consistent with flow cytometry results.

### Drug loaded ICAM-targeted liposomes reduce brain edema

Brain injection of TNF-α leads to reproducible brain edema, as assessed by measuring extravasation of radiolabeled albumin. Liposomes were loaded with dexamethasone (**Supplemental Table 6 and Supplemental Figure 11**) and free dexamethasone, dexamethasone-loaded IgG liposomes, and dexamethasone-loaded ICAM-targeted liposomes were assessed for effects on brain edema (**Figure 5a**, **Supplemental Figure 12**). No significant effects were detected for IV injection of 0.5 mg/kg free dexamethasone (−0.531 ± 26.3% protection) or dexamethasone-loaded IgG liposomes with equivalent drug dose (24.3 ± 18.9% protection). Dexamethasone-loaded ICAM-targeted liposomes provided near complete protection from edema (88.5 ± 14.6% protection) (**Figure 5b**). Precluding effects of the liposomes themselves, neither empty IgG liposomes (40.7 ± 23.5% protection) nor ICAM-targeted liposomes (−4.14 ± 29.81% protection) provided significant protection against edema (**Figure 5b**).

**Figure 5.**
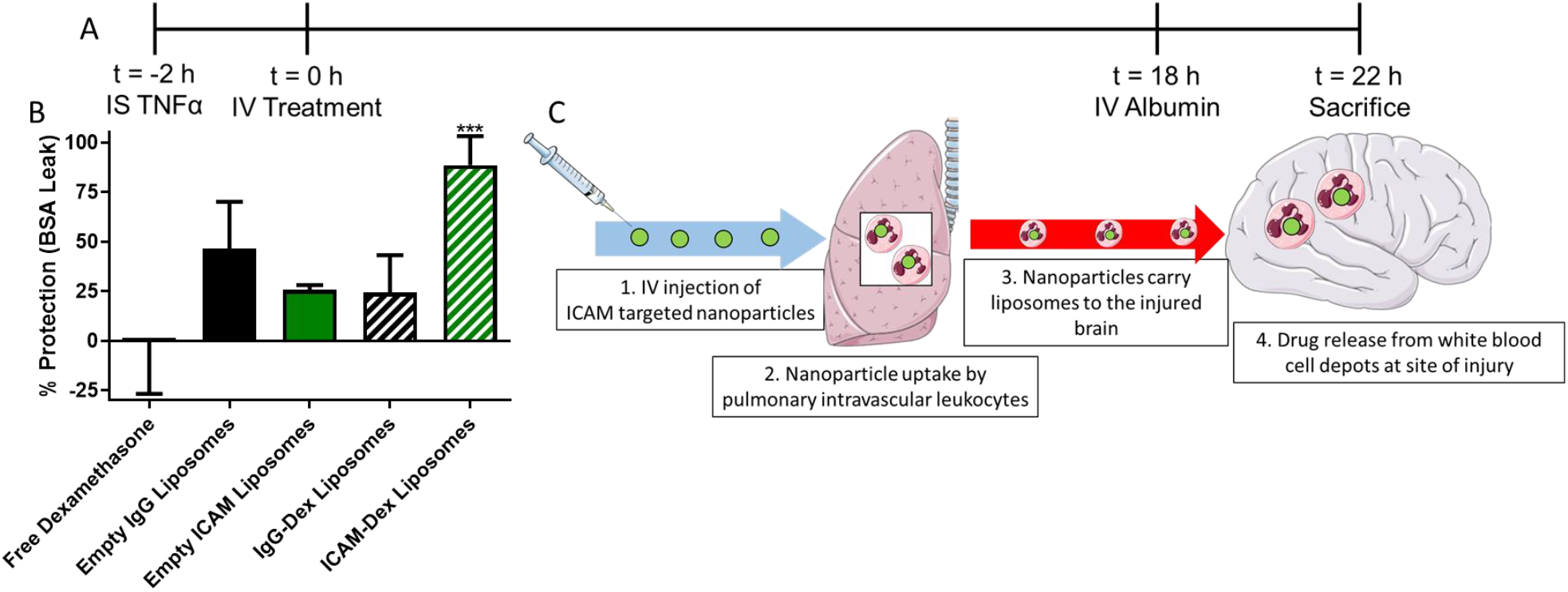
ICAM-targeted dexamethasone (Dex) liposomes protect mice from TNF-induced brain edema. A) Experimental timeline. B) Protective effects of ICAM-targeted dexamethasone liposomes (0.5 mg/kg dexamethasone). As controls for Dex-loaded liposomes, equivalent doses of empty IgG or ICAM-targeted liposomes were tested. % protection was calculated assuming 100% protection as equivalent to edema induced by sham injury and 0% protection as equivalent to edema induced by TNF injury without treatment (**Supplemental Figure 12**). C) Proposed model of leukocyte-mediated drug delivery. Data displayed as mean ± SEM. Comparisons made by 1-way ANOVA with Dunnett’s post-hoc test vs. untreated (solid line, 0% protection). N ≥ 3/group.

Complete blood counts were performed to assess the impact of dexamethasone loaded into ICAM-targeted liposomes and other formulations on blood cells (**Supplemental Figure 13**). ICAM-targeted dexamethasone liposomes led to a reduction in lymphocytes, consistent with the known mechanism of action of the drug, but no other blood cell parameters were affected by treatment, indicating that the therapeutic effect of ICAM-targeted liposomal dexamethasone represents localized action in the brain rather than a systemic effect.

## DISCUSSION

Development of effective therapies for neurological disorders presents formidable challenges including limited success in targeted drug delivery to the brain and especially into the required components of the parenchyma – neurons, glia, etc. The pressing need for effective targeted therapies is especially aggravated in patients suffering from acute brain injuries including stroke, traumatic brain injury, neuroinflammation, and intracranial hemorrhage. These patients present additional challenges for the pharmacotherapy including but not limited to complicating factors, including: 1) rapid disease progression, 2) multiple pathophysiological factors, and 3) poor tolerance for adverse effects.

Harnessing natural host defense mechanisms by loading nanoparticles into leukocytes responding to signals emanating from the injured brain is an attractive strategy for drug delivery. In this case, delivery to sites of injury would be controlled by the natural homing mechanisms used by leukocytes to reach the brain (e.g. emanating chemokine gradients, cell adhesion molecules, etc.). Leukocytes have been used as carriers in chronic neurodegenerative conditions following *ex vivo* loading of drugs and reinfusion into animals^21, 42, 43^. In these studies, it was suggested that leukocytes (or leukocyte-derived extracellular vesicles) could not only reach the brain, but also mediate transfer of their cargo into neurons in order to elicit a pharmacologic response.^44, 45^. Through direct targeting of leukocytes *in vivo*, the need for complex *ex vivo* manipulations can be bypassed in a manner that is permissive for selective delivery into the brain parenchyma

We postulated that direct targeting of pulmonary intravascular leukocytes would be a viable strategy to achieve selective drug delivery to injured regions of the brain, which has several potential advantages vs. *ex vivo* modification, including: 1) treatment can be initiated rapidly after injury, without the need for *ex vivo* modification of cells, 2) all leukocytes accessible to IV injected mAb/nanoparticles are potential targets for loading, 3) selection for specific leukocyte phenotypes is possible by targeting to specific markers, and 4) leukocytes could be converted into drug depots/biofactories that concentrate drugs in the inflamed region where their activity is required. By targeting ICAM expressed on the surface of activated leukocytes, a decline in lung concentrations was seen in parallel with delivery to the injured brain.

Following IV administration, affinity ligands directed towards many vascular epitopes have low levels of delivery to organs such as the brain, in part due to first pass binding to the pulmonary endothelium. However, by directly targeting cell populations that transiently reside in the lungs (e.g. intravascular leukocytes), conversion of the lung from a competitor into an active participant in delivery to the brain is feasible. The data presented above show that targeting to molecules expressed on all (CD45) and activated (ICAM) leukocytes permits delivery to the brain, despite significant uptake by the lungs. The purported mechanism for this delivery is that ICAM-targeted nanoparticles rapidly bind to pulmonary intravascular leukocytes and remain associated with leukocytes, likely in an intracellular compartment following CAM-mediated endocytosis^46^, as they migrate to the brain in response to inflammatory signaling.

Following studies aimed at suggesting a mechanism of delivery to the brain, we pursued therapeutic studies to elucidate the therapeutic relevance of this leukocyte-based drug delivery strategy. We selected the small molecule corticosteroid dexamethasone as a therapeutic agent. Notably, dexamethasone has been tested in clinical trials for treatment of acute ischemic stroke, but ultimately failed due to off-target effects. Among its pleiotropic effects, dexamethasone downregulates expression of the following: inducible CAMs, inflammatory cytokine expression (IL-1, IL-6, TNF-α), cyclooxygenase-2, collagenase, and NF-κB^47^. We hypothesized that ICAM-targeted dexamethasone-loaded liposomes would provide selective delivery of dexamethasone to the injured brain. The results presented here demonstrate that IV injection of dexamethasone 2 hours post-TNF injury was only able to prevent brain edema when encapsulated in ICAM-targeted liposomes (**Figure 5b**). These results are likely due to not only changes in local brain concentrations of dexamethasone, but also due to direct effects on leukocytes targeted by this strategy.

In summary, we have developed a novel approach for direct, *in vivo* loading of activated leukocytes with nanoparticles via targeting to ICAM-1 (**Figure 5c**). We propose the following mechanism for brain delivery whereby the pulmonary intravascular leukocytes: 1) respond to signals emanating from the injured brain and change their activation status and local concentration, 2) are targeted by αICAM mAbs and nanoparticles, and 3) shuttle the taken up αICAM mAb/nanoparticles from the lungs to the injured region of the brain. Our results show that direct leukocyte targeting provides a steady accumulation of nanoparticles into the brain parenchyma following induction of acute neurovascular inflammation. Essentially all of the targeted nanoparticles in the brain were associated with leukocytes, namely monocytes/macrophages and neutrophils. Injection of ICAM-targeted, dexamethasone-loaded liposomes into mice two hours post-TNF injury was able to completely protect mice from injury-induced brain edema. By harnessing natural leukocyte migration patterns, this strategy provides enhanced selectivity for the injured region of the brain and has potential for applications in other acute neurovascular inflammatory injuries such as stroke.

## MATERIALS AND METHODS

### Reagents

Reagents for iodination of proteins were obtained from the following sources: Na^125^I (PerkinElmer, Waltham, MA), 1,3,4,6-tetrachloro-3α,6α-diphenyl-glycouril (Iodogen^®^) (Pierce, Rockford, IL). Polystyrene beads (190 nm) were purchased from Bangslabs (Fishers, IN). All lipids for liposome formulation were obtained from Avanti Polar Lipids (Alabaster, AL). Pooled rat IgG (rIgG) was purchased from Invitrogen (Carlsbad, CA). All other chemicals and reagents were purchased from SigmaAldrich (St. Louis, MO), unless specifically noted.

### Animals

All animal studies were carried out in accordance with the Guide for the Care and Use of Laboratory Animals (National Institutes of Health, Bethesda, MD) and all animal protocols were approved by the University of Pennsylvania Institutional Animal Care and Use Committee. All animal experiments were carried out using male, 6-8 week old C57BL/6 mice (20-25 g) (The Jackson Laboratory, Bar Harbor, ME).

### Protein Production and Purification

Anti-ICAM mAb (YN1) was produced and purified from hybridoma supernatants, as described previously^48^. Purification of YN1 was performed using Protein G affinity chromatography.

### Radiolabeling

Antibodies (YN1, rIgG) were radiolabeled with ^125^I via the Iodogen^®^ method. Briefly, tubes were coated with 100 μg of Iodogen^®^ reagent were incubated with antibodies (1-2 mg/mL) and Na^125^I (0.25 μCi/μg protein) for 5 minutes on ice. Residual free iodine was removed from the bulk solution using a desalting column and thin layer chromatography was used to confirm the efficiency of radiolabeling. As a quality control step, all proteins were confirmed to have <10% free ^125^I prior to further use.

### Polystyrene Nanoparticle Conjugation

Carboxylated, polystyrene beads were conjugated to antibodies (rIgG, YN1) via reaction of N-hydroxysulfosuccinimide (sulfo-NHS) (0.275 mg/mL), 1-ethyl-3-(3-dimethylaminopropl)carbodiimide HCl (EDC) (0.1 mg/mL), and 200 antibody molecules/bead. For experiments involving radioisotope tracing, 15% of the total antibody added to the reaction was ^125^I-labeled rIgG. NC size and polydispersity index (PDI) were confirmed via dynamic light scattering (DLS).

### Liposome Formulation

Liposomes were prepared as described previously ^48^. Briefly, 1,2-dipalmitoyl-sn-glycero-3-phosphocholine (DPPC), cholesterol, and 1,2-distearoyl-sn-glycero-3-phosphoethanolamine-N-[azido(polyethyleneglycol)-2000 (DSPE-PEG2000-azide) were mixed in a molar ratio of 54:40:6. Liposomes were prepared via the thin film extrusion method. To form drug loaded liposomes, the lipid film was hydrated in a solution containing 20 mg/mL of dexamethasone-21-phosphate in phosphate buffered saline (PBS), pH 7.4. The resulting vesicles were extruded through 200 nm polycarbonate membranes.

### Lipid Nanoparticle (LNP) Formulation

LNPs were prepared via microfluidic mixing as previously described^49^. Briefly, an ethanol phase was prepared by combining ionizable lipid, 1,2-dioleoyl-sn-glycero-3-phosphoethanolamine (DOPE), cholesterol, 1,2-dimyristoyl-*sn*-glycero-3-phosphoethanolamine-*N*-[methoxy(polyethylene glycol)-2000] (C14-PEG2000) at molar ratios of 35:16:46.5:2.5, respectively. Separately, an aqueous phase was prepared by resuspending scrambled siRNA sequences in 10 mM citrate buffer to a concentration of .75 mg/mL. Ethanol and aqueous phases were then mixed in a single channel microfluidic device at a 3:1 ratio using a syringe pump^50^. LNPs were dialyzed against 1x PBS for 2 hours at room temperature, followed by sterile filtration using .22 μm syringe filters.

Conjugation of antibodies to the liposome surface was carried out using strain-promoted alkyne-azide cycloaddition. Antibodies were functionalized by reacting with a 5-fold molar excess of dibenzocyclooctyne-PEG_4_-NHS ester (DBCO-PEG4-NHS) (Click Chemistry Tools, Scottsdale, AZ) for 30 minutes at room temperature. Unreacted DBCO-PEG4-NHS was removed via centrifugation through a molecular weight cutoff (MWCO) filter. Liposomes were conjugated with DBCO-functionalized antibodies by reacting for 4 hours at 37 °C. For experiments involving radiotracing, 10% of the total antibody added was ^125^I-labeled rIgG. Unconjugated antibody was removed from the liposomes using gel filtration chromatography. The size, distribution, and concentration of liposomes was determined using DLS and nanoparticle tracking analysis (Malvern Panalytical, Westborough, MA).

### Dexamethasone Loading and Release

Both the amount of dexamethasone loading into liposomes and kinetics of release were assessed using reverse phase high performance liquid chromatography (HPLC). The mobile phase consisted of 30% v/v acetonitrile, 70% v/v water, and 0.1% v/v trifluoroacetic acid. Buffer was run at a flow rate of 0.6 mL/minute through a C8 column (Exlipse XDB-C8, 3 μm, 3.0×100 mm, Phenomenex). Dexamethasone was detected using UV absorbance at 240 nm and the assay had a linear range of 1.56 – 100 μg/mL (**Supplemental Figure 14**). Drug release was measured by dialyzing loaded particles against a large excess of PBS, pH 7.4 at 37 °C and collecting samples at designated time points.

### TNF Injury Model

Neurovascular inflammation was induced in mice via a unilateral injection of TNF-α (0.5 μg/mouse, 2.5 μL, BioLegend) into the striatum using a stereotaxic frame at the following coordinates relative to the bregma: 0.5 mm anterior, 2.0 mm lateral, −3 mm ventral^10^. At different times relative to TNF-α injection (1-24 hours), mice were injected intravenously with a bolus dose of either mAbs (5 μg) or nanoparticles (polystyrene beads, liposomes). Animals were perfused with 20 mL of PBS, pH 7.4 prior to collecting organs for further analysis. For pharmacokinetic and biodistribution studies, the amount of radioactivity in blood and organs was measured using a gamma counter (Wizard2, PerkinElmer, Waltham, MA).

### Transmission Electron Microscopy

Visualization of NC uptake in the lungs shortly after injection was performed using TEM, as previously described^51^. Briefly, 30 minutes post-injection, lungs were fixed with 2.5% glutaraldehyde and 4% paraformaldehyde in 0.1 M sodium cacodylate buffer, then processed into 80-90 nm-thin resin-embedded sections to visualization by TEM.

### Intravital Microscopy

After removing the meninges, a cranial window was opened in one parietal bone of mice. This window was sealed with a glass coverslip and a cannula (PlasticsOne, Roanoke, VA) was placed into the subarachnoid space adjacent to the window (1 mm depth). Animals were allowed to recover for 5 days between opening of the cranial window and injection of TNF-α to prevent any artifacts related to surgery-induced inflammation. *In vivo* imaging was performed in real time with a Stereo Discovery V20 fluorescence microscope (Carl Zeiss AG, Oberkochen, Germany).

### Flow Cytometry

Single cell suspensions of brain were produced as described previously ^9, 52^. Briefly, tissues were enzymatically digested with dispase and collagenase for 1 hour at 37 °C, followed by addition of 600 U/mL DNase Grade II. Tissue digests were demyelinated in Percoll and ACK buffer (Quality Biological, Gaithersburg, MD) was added to lyse any residual RBCs. Samples were then filtered through: 1) 100 μm nylon strainers and 2) 70 μm nylon strainers (ThermoFisher).

Cells were then stained with appropriate antibodies (**Supplemental Table 7**). Briefly, 2×10^6^ cells were labeleld per tube in PBS containing 2% v/v fetal bovine serum (FBS). Fc receptors were blocked using TruStain FcX PLUS (anti-mouse CD16/32, 1:200 dilution) (BioLegend). In pilot experiments to determine localization of NC in leukocytes (CD45^+^) vs. endothelial cells (CD31^+^), flow cytometry was performed using an Accuri C6plus (Benton Dickinson, San Jose, CA). Detailed subtyping of white blood cells in the brain was performed using the strategy described by Posel et al. using a BD LSRFortessa (Benton Dickinson, San Jose, CA) flow cytometer. Live/dead staining was performed using LIVE/DEAD Fixable Aqua Dead Cell Stain Kit (1:1000 dilution, ThermoFisher). In this assay uptake by the following cell types was defined: 1) microglia (CD45^mid^), T-cells (CD45^hi^CD3^+^), neutrophils (CD45^hi^Ly6G^+^), monocytes/macrophages (CD45^hi^CD3^-^Ly6G^-^CD11b^+^Ly6C^+^). Analysis of flow cytometry data was performed using the BD Accuri C6 software (Benton Dickinson, San Jose, CA) and FlowJo v10.6.2 (Tree Star).

### Histology

TNF brains injected with IgG- or αICAM conjugated NCs were perfused, harvested 24 hours-post injected, and fixed in 4% paraformaldehyde. After freezing in tissue freezing medium, the brains were sectioned at 20 μm thickness. Tissue sections were then permeabilized and blocked in blocking solution (5% normal goat serum and 0.3% Triton X-100 in PBS) for 1 hour at room temperature, then incubated overnight at 4 °C with primary antibodies (**Supplemental Table 8**) in blocking solution. After washing with PBS, the sections were incubated with secondary antibodies conjugated with Alexa fluorophores (1:200, Invitrogen) in PBS for 1 hour at room temperature. After washing, the sections were counterstained with nuclei dye 4’-6-Diamidino-2-phenylindole (DAPI, Southern, Biotech). The images were taken by Leica DM6000 Widefield Microscope.

### Therapeutic Studies

The effects of dexamethasone on TNF-induced brain edema were assessed as described in our previous publication^9^. Briefly, 2 hours post-TNF injection, mice were dosed IV with either: 1) 0.5 mg/kg dexamethasone, 2) empty liposomes (either αICAM or IgG coated), or 3) 0.5 mg/kg liposomal dexamethasone (either αICAM or IgG coated). 20 hours after TNF injection, mice were injected with ^125^I-labeled bovine serum albumin (BSA, ~3×10^6^ cpm/mouse), which was then allowed to circulate for 4 hours. After BSA circulation, mice were perfused with 20 mL of PBS, pH 7.4, over 5 minutes and organs were harvested. Edema was determined by measuring the relative concentration of extravasated BSA in the brain to the concentration in the bloodstream. For calculations of therapeutic efficacy, 0% protection was defined using PBS-treated, TNF-injured mice and 100% protection was defined using PBS-treated, sham-injured mice.

### Complete Blood Counts

At designated time points, blood was collected from mice into tubes containing EDTA. Blood cells were analyzed using an Abaxis VetScan HM5 Hematology Analyzer and all values were normalized to the mean value obtained for naïve mice.

### Statistics

All statistical tests were performed using GraphPad Prism 8 (GraphPad Software, San Diego, CA). * denotes p<0.05, ** denotes p<0.01, *** denotes p<0.001, **** denotes p<0.0001.

