## Supplemental Data for "Targeted nanocarriers coopting pulmonary leukocytes for drug delivery to the injured brain"

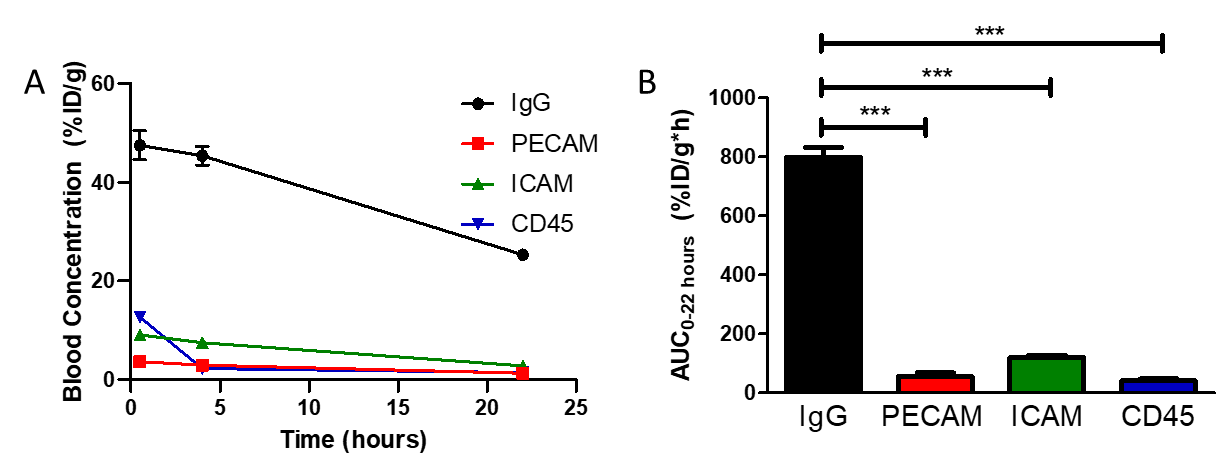

**Supplemental Figure 1**. Blood pharmacokinetics of mAbs against distinct vascular accessible epitopes following IV injection 2 hours post-TNF-α injury. A) Blood concentration vs. time data. B) Area under the concentration vs. time curve from 0 – 22 hours (AUC_0-22 h_). Data represented as mean ± SEM. *** denotes p<0.001 by 1-way ANOVA with Dunnett’s post-hoc test. N = 3/group.

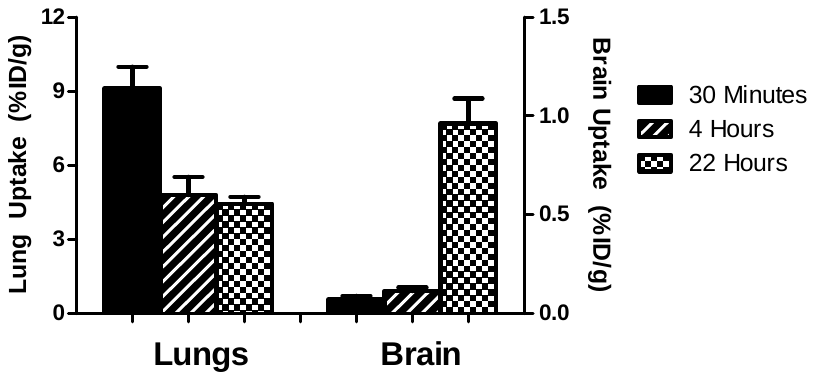

**Supplemental Figure 2.** Lung and brain pharmacokinetics of control IgG injected 2 hours post-TNF-α injury. Experimental timeline as in Figure 2b. Data represented as mean ± SEM. N = 3/group.

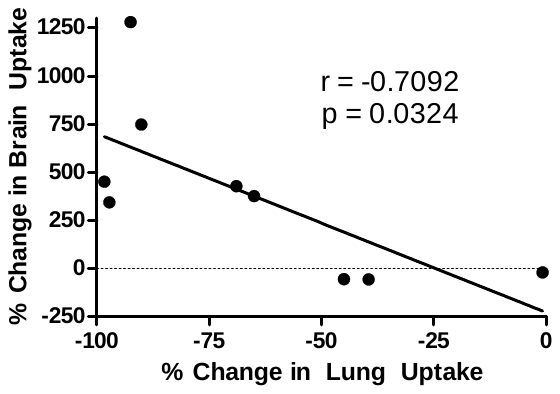

**Supplemental Figure 3**. Changes in brain uptake correlate with clearance from the lung. Data from Figure 2 displayed as individual animals. Pearson’s correlation analysis was used to derive the correlation coefficient (r).

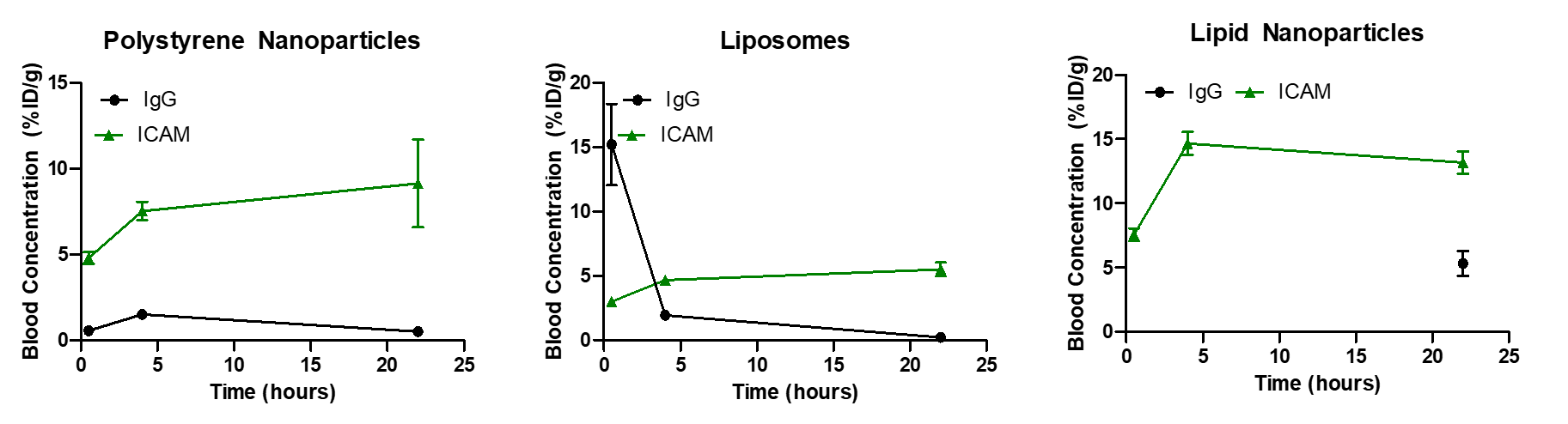

**Supplemental Figure 4.** Blood pharmacokinetics of nanoparticles injected intravenously 2 hours post-TNF-α injury.

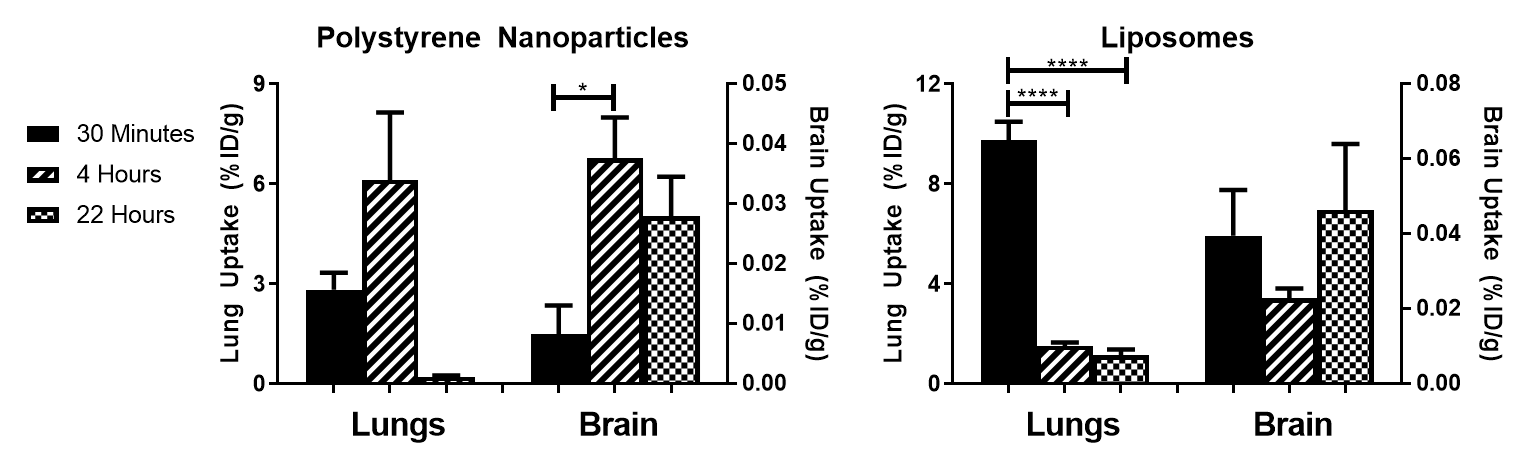

**Supplemental Figure 5.** Lung and brain pharmacokinetics of control IgG injected 2 hours post-TNF-α injury. Experimental timeline as in Figure 3a. Comparisons made by 1-way ANOVA with Dunnett’s post-hoc test vs. 30 minutes. Data represented as mean ± SEM. N = 3/group.

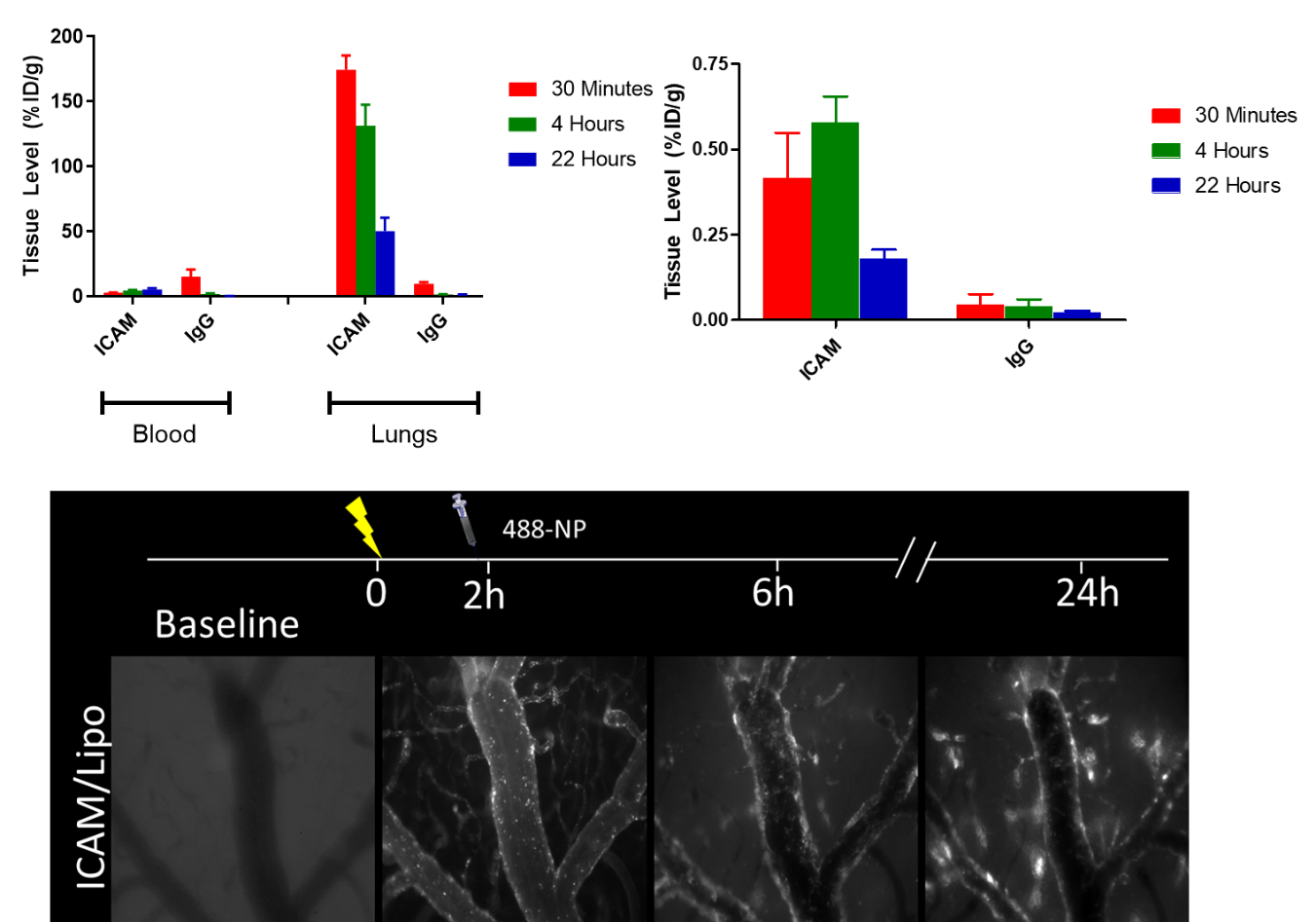

**Supplemental Figure 6**. Cranial window intravital microscopy of ICAM-targeted liposomes in the brain at designated time points after TNF-α injury.

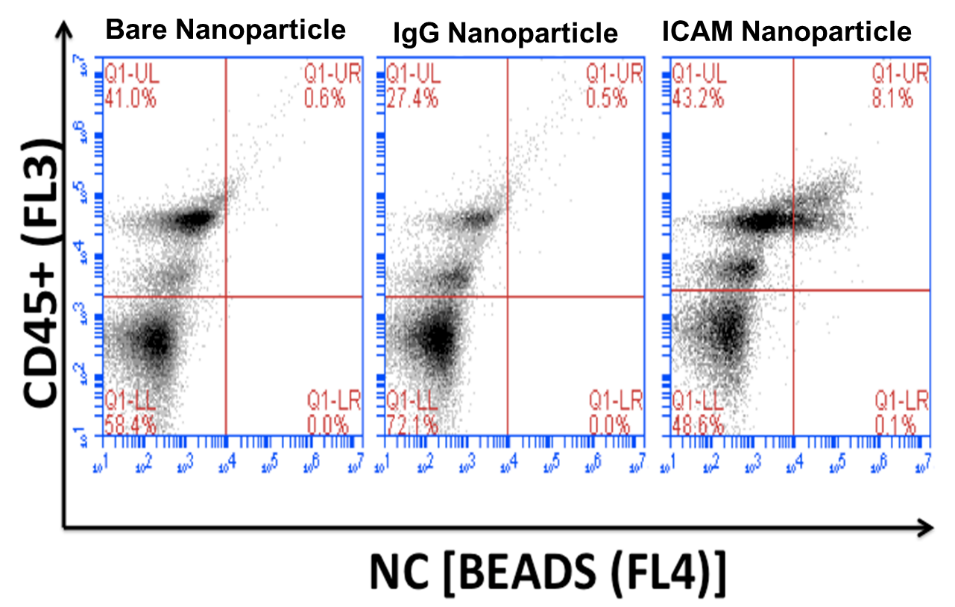

**Supplemental Figure 7.** Representative flow cytometry dot-plots of polystyrene bead (bare, IgG, ICAM) cellular uptake in the brain 22 hours post-injection (24 hours post-TNF).

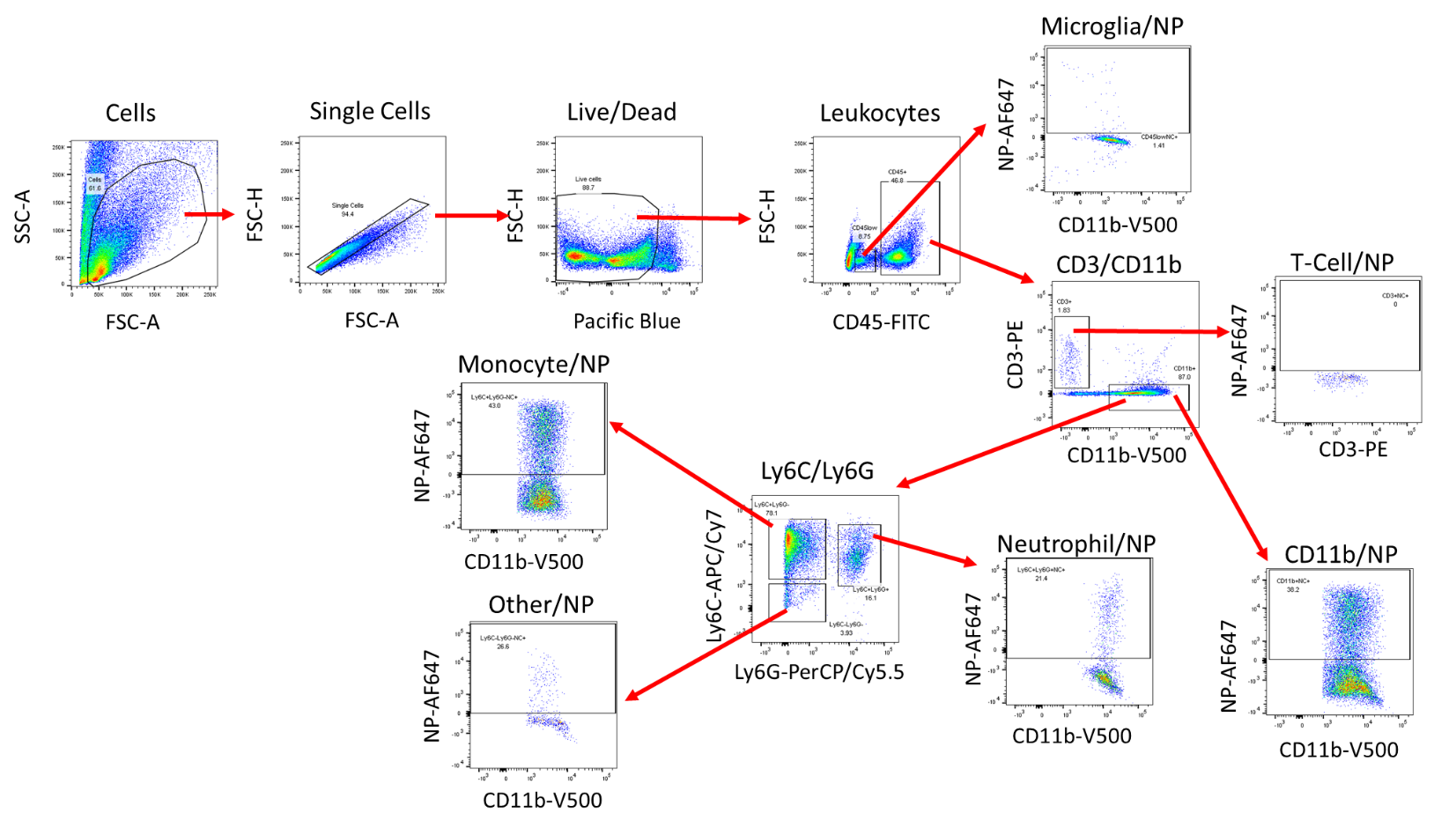

**Supplemental Figure 8**. Gating scheme for detailed flow cytometry of leukocyte sub-types in brain single cell suspensions.

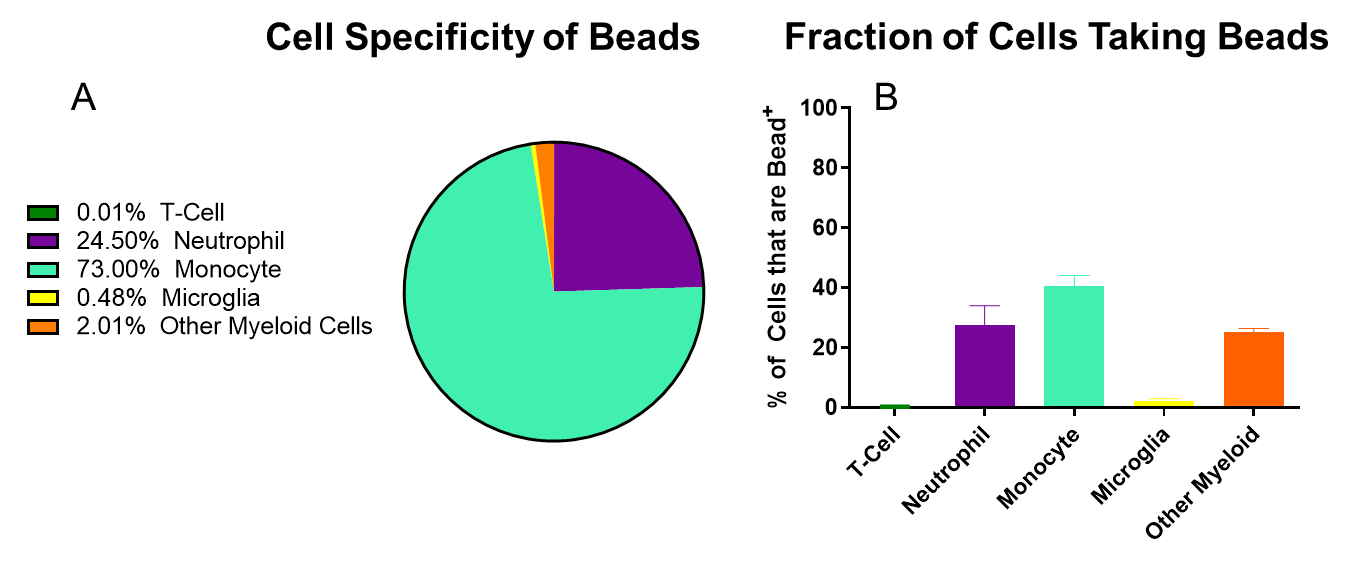

**Supplemental Figure 9.** Flow cytometry of ICAM-targeted polystyrene nanoparticle distribution in leukocytes in the brain 22 hours post-injection (24 hours post-TNF). A) Fraction of nanoparticle-positive leukocytes for each sub-type. B) Fraction of recovered cells that were nanoparticle^+^. T-Cell: CD45^+^CD11b^-^CD3^+^, Neutrophil: CD45^+^CD11b^+^Ly6G^+^, Monocyte: CD45^+^CD11b^+^Ly6C^+^Ly6G^-^, Microglia: CD45^mid^, Other Myeloid: CD45^+^CD11b^+^Ly6C^-^Ly6G^-^. Data represented as mean ± SEM. N = 3/group

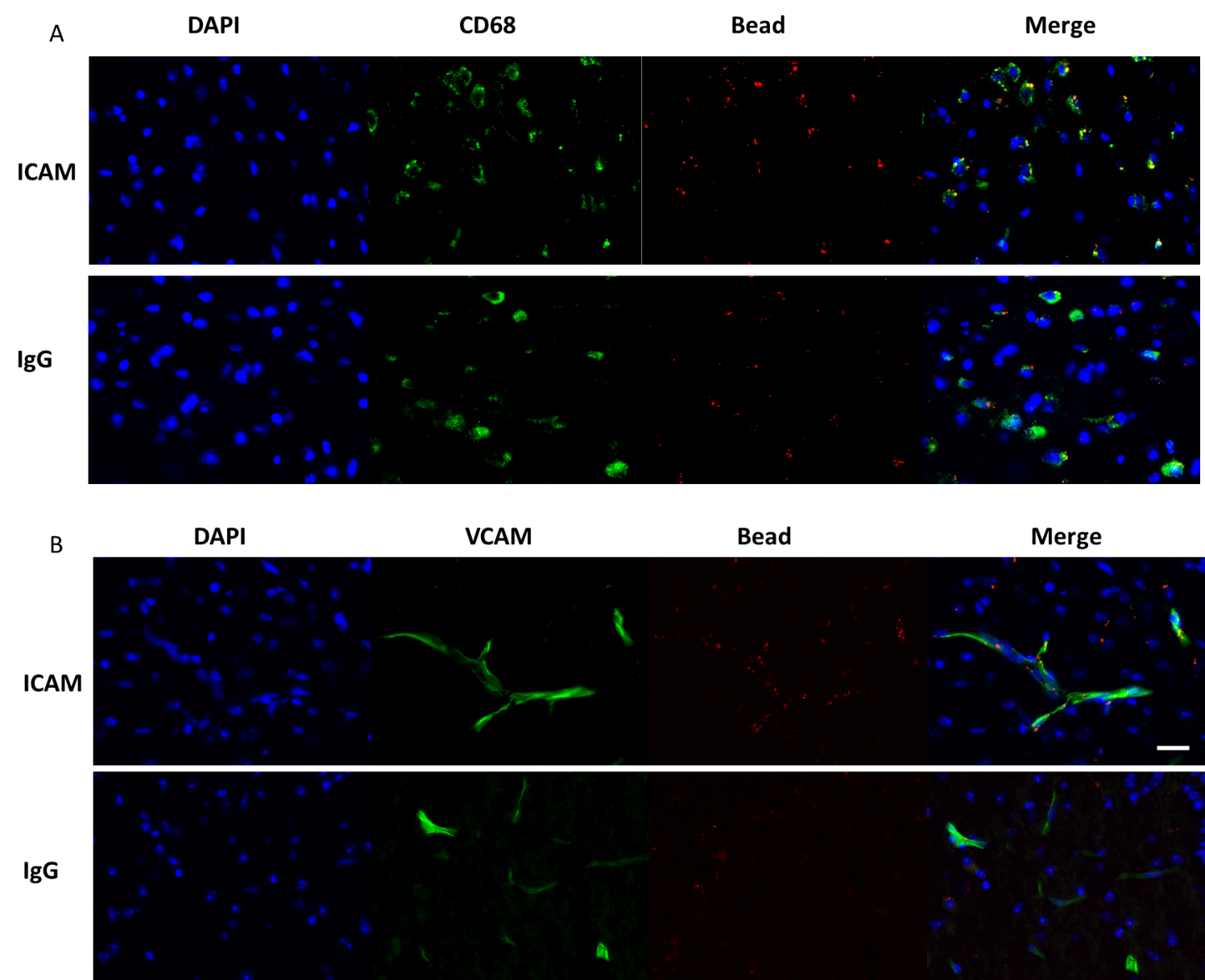

**Supplemental Figure 10:** Histology of brain slices taken 22 hours post-injection of ICAM-targeted or IgG polystyrene nanoparticles (24 hours post TNF-α injection). Staining was performed to determine co-localization with A) macrophages (CD68) and B) endothelial cells (VCAM). Scale bar: 50 μm.

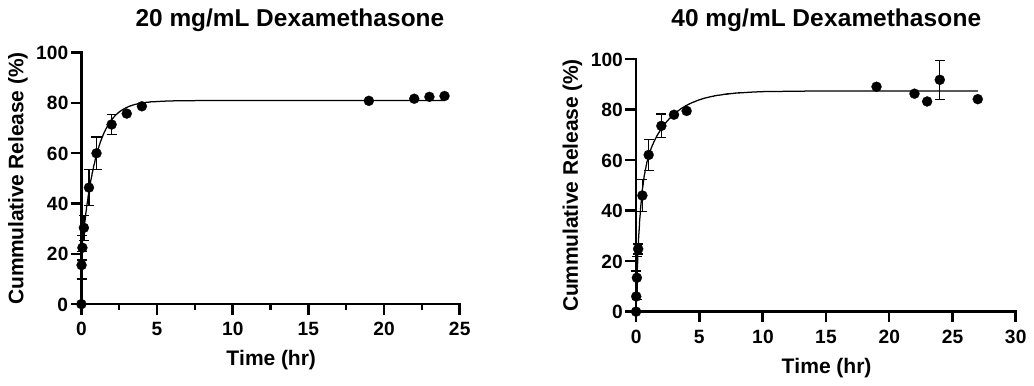

**Supplemental Figure 11**. Dexamethasone release from liposomes incubated in PBS at 37°C.

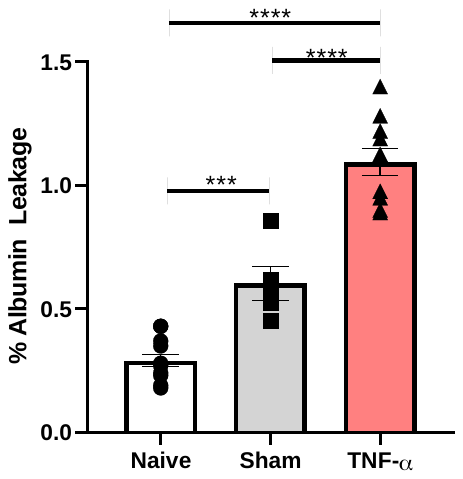

**Supplemental Figure 12**. TNF-α injury induces albumin leak into the brain. 20 hours after injection of TNF-α mice were IV injected with radiolabeled albumin. Following perfusion, the degree of albumin leak was calculated as the percentage of brain albumin concentrations vs. blood. Data represented as mean ± SEM. Comparisons made by 1-way ANOVA with Tukey’s post-hoc test. N = 5-12/group.

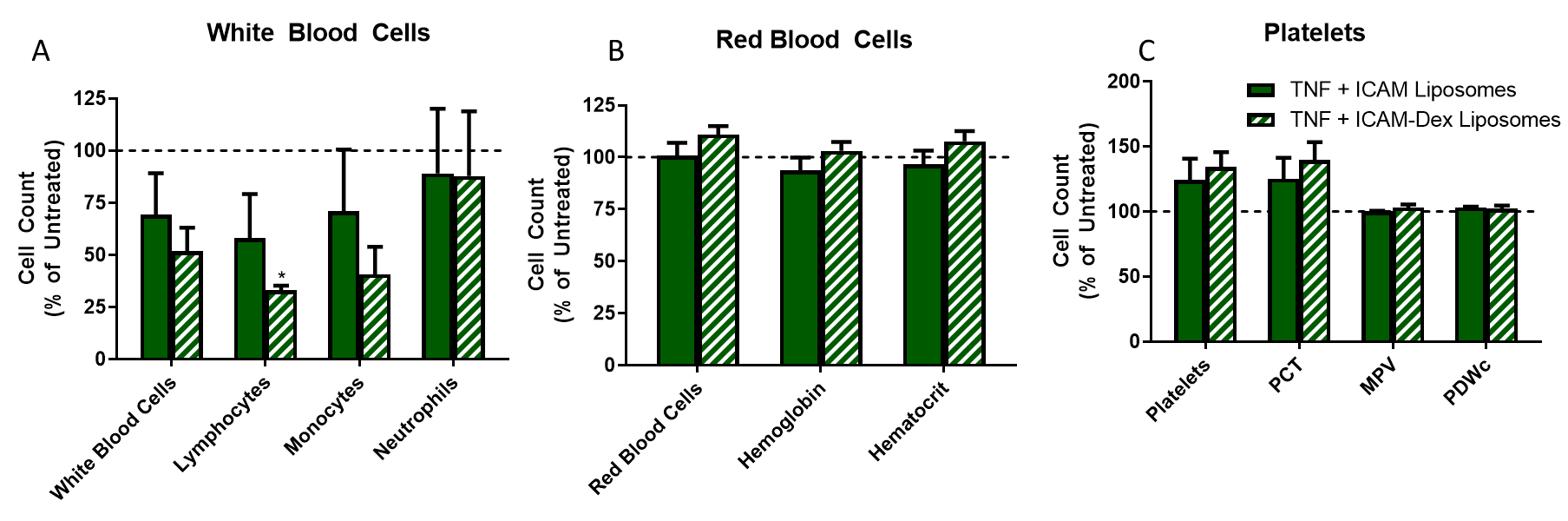

**Supplemental Figure 13**. Complete blood counts 22 hours post-injection of ICAM-targeted liposomes and dexamethasone-loaded ICAM-targeted liposomes (24 hours post-TNF-α). A) White blood cells, B) Red blood cells, C) Platelets. Data represented as mean ± SEM. Dashed line represents values for untreated mice. Comparisons made by 1-way ANOVA with Dunnett’s post-hoc test vs. untreated. N ≥ 3/group.

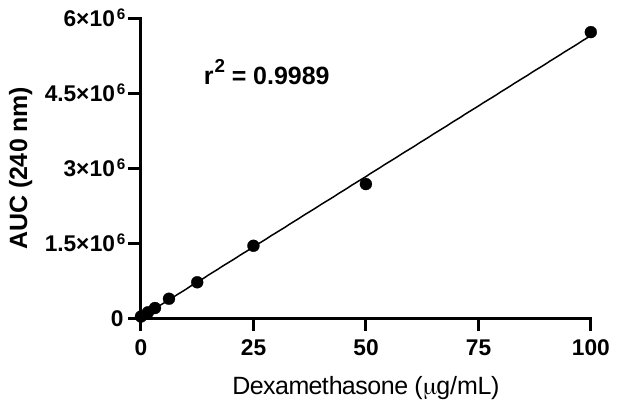

**Supplemental Figure 14**. Representative standard curve for dexamethasone HPLC assay.

| **αICAM** | Blood | Lung | | Liver | | | Kidney | | Heart | | | | Spleen | Brain | | |
| --- | --- | --- | --- | --- | --- | --- | --- | --- | --- | --- | --- | --- | --- | --- | --- | --- |
| Naïve | 4.69±1.59 | | 106±13 | | 19.6±2.1 | | 9.51±0.90 | | | 3.53±0.29 | | 47.8±4.1 | | | | 0.347±0.049 |
| 1 hour | 6.04±0.21 | | 216±8** | | 17.5±1.0 | | 29.2±1.3*** | | | 10.2±0.3*** | | 24.3±2.4** | | | | 0.731±0.049 |
| 2 hours | 7.23±0.22 | | 258±16*** | | 33.2±4.8** | | 19.3±0.8*** | | | 6.40±0.61** | | 44.5±4.2 | | | | 0.655±0.162 |
| 3 hours | 5.99±0.59 | | 194±16** | | 18.7±0.6 | | 31.0±1.3*** | | | 11.9±0.5*** | | 31.6±1.0* | | | | 0.882±0.028 |
| 6 hours | 3.19±0.42 | | 115±12 | | 16.8±0.6 | | 11.1±1.0 | | | 5.86±0.40* | | 22.7±1.8*** | | | | 1.31±0.16** |
| 24 hours | 5.02±0.13 | | 108±24 | | 14.2±1.0 | | 7.74±1.97 | | | 4.79±0.86 | | 27.2±4.6** | | | | 2.43±0.24*** |
| **αCD45** | Blood | | Lung | | Liver | Kidney | | Heart | | | Spleen | | | | Brain | |
| Naïve | 12.7±0.7 | | 13.2±2.2 | | 27.8±2.8 | 5.01±0.13 | | 1.77±0.22 | | | 189±16 | | | | 0.034±0.005 | |
| 2 hours | 12.7±0.6 | | 60.5±10.2** | | 33.5±0.3 | 5.68±0.37 | | 0.946±0.110 | | | 136±17 | | | | 0.061±0.010 | |
| 24 hours | 12.6±0.6 | | 35.9±7.3 | | 27.4±1.2 | 7.30±0.70* | | 1.56±0.29 | | | 215±16 | | | | 0.536±0.042**** | |
| **IgG** | Blood | | Lung | | Liver | Kidney | | Heart | | | Spleen | | | | Brain | |
| Naïve | 30.4±0.9 | | 0.931±0.526 | | 3.69±0.52 | 4.29±0.98 | | 1.28±0.17 | | | 2.36±0.01 | | | | 0.077±0.022 | |
| 1 hour | 50.2±1.9** | | 1.33±0.42 | | 15.8±2.4*** | 11.7±2.2 | | 1.26±0.25 | | | 9.75±0.51 | | | | 0.073±0.007 | |
| 2 hours | 41.5±2.6 | | 6.88±2.90* | | 11.2±1.3** | 2.11±0.09 | | 1.58±0.31 | | | 7.33±0.62 | | | | 0.033±0.006 | |
| 3 hours | 49.0±0.5** | | 6.30±1.31 | | 16.2±1.0*** | 16.4±5.7* | | 1.26±0.16 | | | 9.90±1.85 | | | | 0.074±0.025 | |
| 6 hours | 31.0±5.6 | | 0.862±0.060 | | 4.12±0.86 | 4.30±0.35 | | 1.33±0.26 | | | 3.70±0.84 | | | | 0.069±0.021 | |
| 24 hours | 43.2±2.1* | | 0.917±0.293 | | 4.65±0.02 | 4.09±0.11 | | 1.37±0.21 | | | 4.33±0.21 | | | | 0.109±0.021 | |

**Supplemental Table 1**. Biodistribution of αICAM, αCD45, and IgG injected at different times post-TNF-α injury. Values reported as percent of injected dose/g organ (%ID/g). Samples were collected 30 minutes after IV injection of mAb. Data reported as mean ± SEM. Comparisons made by 1-way ANOVA with Dunnett’s post-hoc test vs. naïve. * denotes p<0.05, ** denotes p<0.01, *** denotes p<0.001, **** denotes p<0.0001. N = 3/group.

| **αPECAM** | Blood | Lung | Liver | Kidney | | Heart | Spleen | Brain |
| --- | --- | --- | --- | --- | --- | --- | --- | --- |
| 30 minutes | 3.64±0.73 | 133±22 | 22.6±4.2 | 26.0±5.0 | | 19.6±4.7 | 32.3±6.5 | 3.99±0.61 |
| 4 hours | 2.91±0.44 | 143±16 | 26.0±4.9 | 30.7±4.3 | | 22.4±4.5 | 37.2±9.1 | 3.42±0.47 |
| 22 hours | 1.27±0.22 | 90.4±5.1 | 19.8±0.4 | 20.6±1.3 | | 15.0±1.0 | 18.8±2.2 | 2.27±0.08 |
| **αICAM** | Blood | Lung | Liver | Kidney | | Heart | Spleen | Brain |
| 30 minutes | 9.08±0.29 | 223±28 | 23.1±2.6 | 30.4±2.0 | | 11.2±1.4 | 38.1±9.4 | 0.605±0.053 |
| 4 hours | 7.49±0.35 | 119±10 | 36.8±1.3 | 18.1±1.7 | | 7.14±0.38 | 73.8±2.9 | 0.868±0.035 |
| 22 hours | 2.77±0.37 | 59.3±6.2 | 16.2±0.8 | 7.25±0.32 | | 3.26±0.20 | 29.3±0.4 | 2.52±0.14 |
| **αCD45** | Blood | Lung | Liver | | Kidney | Heart | Spleen | Brain |
| 30 minutes | 12.7 ± 0.6 | 60.5 ± 10.2 | 35.5 ± 1.7 | | 5.68 ± 0.37 | 0.946 ± 0.110 | 136 ± 17 | 0.0610 ± 0.010 |
| 4 hours | 2.22±0.18 | 8.45±0.69 | 23.8±3.7 | | 2.00±0.49 | 0.243±0.023 | 153±36 | 0.146±0.040 |
| 22 hours | 1.42±0.45 | 3.70±0.88 | 3.74±0.49 | | 0.974±0.242 | 0.268±0.084 | 89.6±29.2 | 0.437±0.116 |
| **IgG** | Blood | Lung | Liver | | Kidney | Heart | Spleen | Brain |
| 30 minutes | 47.5±3.0 | 9.12±0.88 | 18.6±0.3 | | 2.67±0.26 | 3.46±0.52 | 11.6±1.7 | 0.071±0.018 |
| 4 hours | 45.4±1.9 | 4.80±0.73 | 6.49±0.90 | | 3.88±0.28 | 2.86±0.11 | 8.49±0.36 | 0.114±0.017 |
| 22 hours | 25.4±0.9 | 4.45±0.28 | 1.65±0.19 | | 1.95±0.35 | 2.44±0.06 | 4.46±0.18 | 0.960±0.125 |

**Supplemental Table 2**. Pharmacokinetics and biodistribution of mAb against endothelial and leukocyte epitopes. Data represented as mean ± SEM. N = 3/group

| **αICAM** | Blood | Lung | | Liver | | Kidney | | Heart | | Spleen | | Brain | |
| --- | --- | --- | --- | --- | --- | --- | --- | --- | --- | --- | --- | --- | --- |
| 30 minutes | 1.91±0.14 | 147±1 | | 25.9±0.5 | | 2.78±0.20 | | 0.578±0.110 | | 21.4±4.4 | | 0.051±0.005 | |
| 4 hours | 2.64±0.53 | 8.82±2.75 | | 14.1±1.1 | | 1.23±0.10 | | 0.341±0.067 | | 29.2±5.1 | | 0.086±0.009 | |
| 22 hours | 3.65±1.02 | 20.2±5.9 | | 8.86±0.94 | | 2.14±0.49 | | 0.726±0.288 | | 25.1±2.6 | | 0.254±0.029 | |
| **IgG** | Blood | | Lung | | Liver | | Kidney | | Heart | | Spleen | | Brain |
| 30 minutes | 0.545±0.066 | | 2.83±0.51 | | 64.7±1.8 | | 0.282±0.025 | | 0.062±0.017 | | 102±7 | | 0.008±0.005 |
| 4 hours | 1.49±0.07 | | 6.12±2.03 | | 41.3±4.3 | | 0.816±0.065 | | 0.215±0.014 | | 67.4±11.2 | | 0.038±0.007 |
| 22 hours | 0.499±0.081 | | 0.222±0.037 | | 16.2±1.0 | | 0.273±0.055 | | 0.053±0.006 | | 16.6±2.7 | | 0.028±0.007 |

**Supplemental Table 3**. Pharmacokinetics and biodistribution of ICAM-targeted and IgG polystyrene nanoparticles injected 2 hours after intrastriatal TNF-α. Experimental design as in **Figure 3a**. Values reported as percent of injected dose/g organ (%ID/g). Samples were collected 30 minutes after IV injection of polystyrene nanoparticles. Data reported as mean ± SEM. N = 3/group.

| **αICAM** | Blood | Lung | | Liver | | Kidney | | Heart | | Spleen | | Brain | |
| --- | --- | --- | --- | --- | --- | --- | --- | --- | --- | --- | --- | --- | --- |
| 30 minutes | 2.99±0.05 | 174±6 | | 36.7±0.4 | | 22.1±1.0 | | 7.99±0.40 | | 40.8±6.7 | | 0.498±0.054 | |
| 4 hours | 4.65±0.25 | 131±9 | | 30.4±1.1 | | 18.3±1.9 | | 6.79±0.39 | | 38.6±6.3 | | 0.702±0.085 | |
| 22 hours | 5.49±0.53 | 50.2±6.0 | | 14.3±1.4 | | 13.2±0.6 | | 4.44±0.17 | | 10.6±0.6 | | 0.287±0.006 | |
| **IgG** | Blood | | Lung | | Liver | | Kidney | | Heart | | Spleen | | Brain |
| 30 minutes | 15.2±3.2 | | 9.76±0.74 | | 56.1±2.4 | | 0.547±0.020 | | 0.367±0.068 | | 65.5±6.2 | | 0.040±0.012 |
| 4 hours | 1.92±0.26 | | 1.52±0.15 | | 46.9±1.5 | | 1.47±0.11 | | 0.101±0.010 | | 49.9±2.5 | | 0.023±0.003 |
| 22 hours | 0.206±0.091 | | 1.16±0.22 | | 32.5±4.9 | | 1.41±0.34 | | 0.090±0.033 | | 25.4±3.4 | | 0.046±0.018 |

**Supplemental Table 4**. Pharmacokinetics and biodistribution of ICAM-targeted and IgG liposomes injected 2 hours after intrastriatal TNF-α =. Experimental design as in **Figure 3a**. Values reported as percent of injected dose/g organ (%ID/g). Samples were collected 30 minutes after IV injection of liposomes. Data reported as mean ± SEM. N = 3/group.

| **αICAM** | Blood | Lung | | Liver | | Kidney | | Heart | | Spleen | | Brain | |
| --- | --- | --- | --- | --- | --- | --- | --- | --- | --- | --- | --- | --- | --- |
| 30 minutes | 7.55±0.49 | 123±9 | | 39.2±1.6 | | 10.7±0.6 | | 5.11±0.32 | | 56.9±7.0 | | 0.281±0.005 | |
| 4 hours | 14.7±0.9 | 53.2±1.4 | | 22.1±0.7 | | 9.89±0.65 | | 2.52±0.25 | | 24.2±3.4 | | 0.313±0.019 | |
| 22 hours | 13.2±0.9 | 14.4±1.3 | | 7.72±0.70 | | 5.66±0.61 | | 2.20±0.36 | | 9.96±0.73 | | 0.344±0.004 | |
| **IgG** | Blood | | Lung | | Liver | | Kidney | | Heart | | Spleen | | Brain |
| 22 hours | 5.31±0.98 | | 0.738±0.243 | | 1.73±0.20 | | 1.81±0.46 | | 0.373±0.053 | | 6.81±0.21 | | 0.0632±0.0459 |

**Supplemental Table 5**. Pharmacokinetics and biodistribution of ICAM-targeted and IgG lipid nanoparticles (LNP) injected 2 hours after intrastriatal TNF-α. Experimental design as in **Figure 3a**. Values reported as percent of injected dose/g organ (%ID/g). Samples were collected 30 minutes after IV injection of LNP. Data reported as mean ± SEM. N = 3/group.

| **Liposome Coating** | **Size (nm)** | **PDI** | **Drug Loading** |
| --- | --- | --- | --- |
| Bare | 134 ± 3 | 0.113 ± 0.005 | 13.3 ± 0.4% |
| IgG | 144 ± 5 | 0.145 ± 0.008 |  |
| αICAM | 139 ± 3 | 0.133 ± 0.007 |  |

**Supplemental Table 6**. Characterization of liposome size, polydisperse index (PDI) and dexamethasone entrapment efficiency.

| **Surface Marker** | **Clone** | **Fluorophore** |
| --- | --- | --- |
| CD3ϵ | 145-2C11 | PE |
| CD11b | M1/70 | V500 |
| CD45.2 | 104 | FITC |
| Ly6G | 1A8 | PerCP-Cy5.5 |
| Ly6C | AL-21 | APC-Cy7 |

**Supplemental Table 7**. Antibodies used for flow cytometry

| **Surface Marker** | **Working concentration** | **Manufacturer** |
| --- | --- | --- |
| CD68 | 10 μg/ml | Bio-Rad |
| VCAM (MK2.7) | 5 μg/ml | In house production |

**Supplemental Table 8.** Antibodies used for histology
